# The RNA-binding activity of the TRIM-NHL protein NHL-2 is essential for miRNA-mediated gene regulation

**DOI:** 10.1101/2024.02.13.580109

**Authors:** Nasim Saadat, Rhys N. Colson, Acadia L. Grimme, Uri Seroussi, Joshua W. Anderson, Julie M. Claycomb, Matthew C. J. Wilce, Katherine McJunkin, Jacqueline A. Wilce, Peter R. Boag

**Author notes:** These authors contributed equally to this work (Nasim Saadat, Rhys N. Colson).

## Abstract

The conserved TRIM-NHL protein, NHL-2, plays a key role in small RNA pathways in *Caenorhabditis elegans*. NHL-2 has been shown to interact with U-rich RNA through its NHL domain, but the importance to its biological function is unknown. We defined the crystal structure of the NHL domain to 1.4 Å resolution and identified residues that affect affinity for U-rich RNA. Functional analysis of an NHL-2 RNA-binding loss-of-function mutant demonstrated defects in the heterochronic pathway, suggesting that RNA binding is essential for its role in this miRNA pathway. Processing bodies were enlarged in the NHL-2 RNA-binding mutant, suggesting a defect in mRNA decay. We also identified the eIF4E binding protein IFET-1 as a strong synthetic interactor with NHL-2 and the DEAD box RNA helicase CGH-1 (DDX6), linking NHL-2 function to translation repression. We demonstrated that in the absence of NHL-2, there was an enrichment of miRNA transcripts associated with the miRNA pathway Argonaute proteins ALG-2 and ALG-2. We demonstrate that NHL-2 RNA-binding activity is essential for *let-7* family miRNA-mediated translational repression. We conclude that the NHL-2, CGH-1, and IFET-1 regulatory axes work with the core miRISC components to form an effector complex that is required for some, but not all, miRNAs.

## Introduction

MicroRNAs (miRNAs) are small, non-coding RNAs that are essential regulators of cell growth, division, differentiation, and development in metazoans (1). In animals, miRNA usually imperfectly base pair with complementary regions within the 3’ untranslated region (3’ UTR) of target mRNAs, with a predominant interaction occurring through the seed region found at nucleotides 2-8 of the miRNA. This triggers a cascade of gene silencing mechanisms, such as translational repression, deadenylation, decapping, and decay (2). The miRNA-induced Silencing Complex (miRISC) is at the heart of miRNA silencing. The minimal miRISC is composed of miRNA bound by an Argonaute (AGO) protein and can directly bind to target mRNAs; however, this complex is insufficient to carry out miRISC-mediated translational repression (3,4). In *C. elegans*, ALG-1 and ALG-2 are the key AGOs involved in miRNA function and generally function redundantly in somatic tissues (5), whereas ALG-2 is also expressed in the germline (6). Additional proteins bind to AGO to form functional effector complexes, the key of which is the scaffolding protein AIN-1/GW182, which functions as a protein-protein interaction platform that facilitates the complex assembly of multiple ribonucleoprotein complexes (RNPs) that determine mRNA fate (2). A highly conserved interaction with AIN-1 are the PAN2-PAN3 and CCR4-CAF1-NOT deadenylase complexes (7). Recent studies have found that the miRISC is not a homogenous complex, but is made up of a variable array of co-factors that may provide additional layers of control at the miRNA, mRNA, and cell type levels (8–11).

In recent years, an expanding body of research has uncovered pivotal roles of RNA-binding proteins in miRNA biogenesis and function (12). RBP-miRISC interplay is complex, with examples of RBPs that enhance or repress miRISC loading onto mRNAs (13,14). This intricate interplay between RBPs and miRNAs has profound implications for the temporal and spatial control of gene expression during development (12). Understanding the nuances of how RBPs participate in miRNA-mediated gene regulation will provide novel insights into the dynamic processes underlying embryogenesis, tissue differentiation, and homeostasis. NHL-2 is a conserved miRISC co-factor that is required for normal miRNA-mediated gene regulation of heterochronic and sex determination genes in *C. elegans* (15,16) and its orthologs in mice (TRIM32, TRIM71) and flies (Mei-P26, Brat) are also critical regulators of stem cell and developmental programs (17–20). We previously showed that the NHL domain is an RNA-binding domain with a preference for poly U sequences and is required for non-miRNA small RNA pathways in *C. elegans* (21), but the functional requirement of RNA binding is unknown.

In order to further understand the role of RNA binding in NHL-2 function we have made use of structural and biophysical information for the NHL-2 NHL domain as well as functional assays and interaction assays using NHL-2 deletion mutants. Here, we report the high-resolution crystal structure of the NHL domain of NHL-2, identification of key amino acid residues for RNA-binding, and *in vivo* functional analysis of a loss-of-function RNA-binding mutant. We used alae and seam cell development as a sensitive indicator of the let-7 microRNA pathway’s functionality and we determined that the normal activity of NHL-2 relies on the C-terminal RING domain, with the RNA-binding activity being crucial for its function. We also discovered that eIF4E-binding proteins and the translational repressor IFET-1 have a strong synthetic interaction with *nhl-2*, suggesting that translational repression is an important part of NHL-2’s function in the miRISC. Interestingly, the Processing body (P-body) size increased in somatic cells in worms containing a loss-of-function RNA-binding mutant of NHL-2, suggesting a defect in mRNA turnover. The absence of NHL-2 resulted in the enrichment of *let-7* miRNA targets that immunoprecipitated with the Argonaute ALG-1/-2 IPs, suggesting defective mRNA regulation of these target mRNAs. We propose that NHL-2 RNA-binding activity is critical for the effector phase of specific miRISCs and that NHL-2 works together with the DEAD box RNA helicase CGH-1 (DDX6) and the eIF4E-binding protein IFET-1(4E-T) to inhibit translation and recruit or stabilise deadenylation and decapping factors that result in *let-7* family mediated mRNA turnover. This study provides new insights into the role of TRIM-NHL RNA-binding proteins in development and links eIF4E-binding proteins to the formation of mature miRISC effector complexes.

## Results

### Structure of the NHL-2 NHL domain

The NHL domain of NHL-2 has previously been shown to preferentially bind to U-rich RNA over other RNA sequences (22). By analogy to other NHL domains such as those of dmBRAT and drLIN41, that have been structurally characterized bound to RNA, this is likely through interactions at the positively charged surface of a six bladed beta-propeller structure (23,24). To obtain a more accurate understanding of the structure of the NHL-2 NHL domain and the specific amino acids likely to underlie RNA binding, we solved the crystal structure of the domain to 1.4 Å resolution. For this, we generated a GST-tagged construct incorporating ceNHL-2 amino acid residues 729-1010, based on the domain boundaries predicted from alignment with homologous NHL domains. Upon cleavage of the GST-tag and purification to high homogeneity, the protein was subjected to crystal screening in a 96 well sitting drop format, at 37 and 4 °C. The NHL domain formed small crystals in 3.5 M sodium formate 4 °C (condition C1 of an Index screen; Hampton Research), with crystals diffracting to 1.4 Å resolution. Larger crystals were formed with a variation to pH 6.8 in 3.5 M sodium formate, 0.1 M HEPES, which diffracted to 1.4 Å resolution.

The NHL-2 NHL domain structure was solved to 1.4 Å resolution using the dmBRAT NHL structure (PDBID: 1QF7) as the molecular replacement model. The full chain was clear in the density except for the C-terminal amino acid residue 1010 which was not detected. Some sidechains were only partially visible in the density including those of Arg729, Arg788, Arg831, Lys922, Lys943, Lys989. Arg806, Gly807, Glu808 lacked density for their sidechains and had poor density for the main chain. After refinement, the model was obtained with R/Rfree values of 16%/17.6 % and good stereochemistry (Table 1). Figure 1A shows cartoon and surface representations of the ceNHL-2 NHL domain from “top” and “side” perspectives. The 6-bladed beta propeller structure is clearly formed as anticipated, with N-and C-terminal beta strands forming an anti-parallel interaction to form blade 6. Clear differences are apparent when compared to other representative NHL domains, with variation in the beta-sheet lengths and twist in each blade as well as the length and orientation of disordered loops. The dmBRAT NHL domain (pdbid:4ZLR solved to 2.3 Å resolution) and drLIN41 NHL domain (PDBID: 6FQC solved to 1.9 Å resolution) structures in complex with their respective RNAs are shown in Figure 1B for comparison. The ceNHL-2 NHL core (without N and C-terminal loops) shows greater topological similarity to dmBRAT NHL with a backbone r.m.s.d of 6.3 Å compared to LIN41 NHL with a backbone r.m.s.d. of 7.4 Å. In fact, the structure-based structural alignment with LIN41 NHL does not superpose the blades in the same register, with the blade 1 of NHL-2 NHL more readily aligning with blade 2 of LIN41 NHL. This variation of the NHL domains reflects the role of the beta propeller as a scaffold from which loops are presented that create unique functional surfaces.

**Figure 1.**
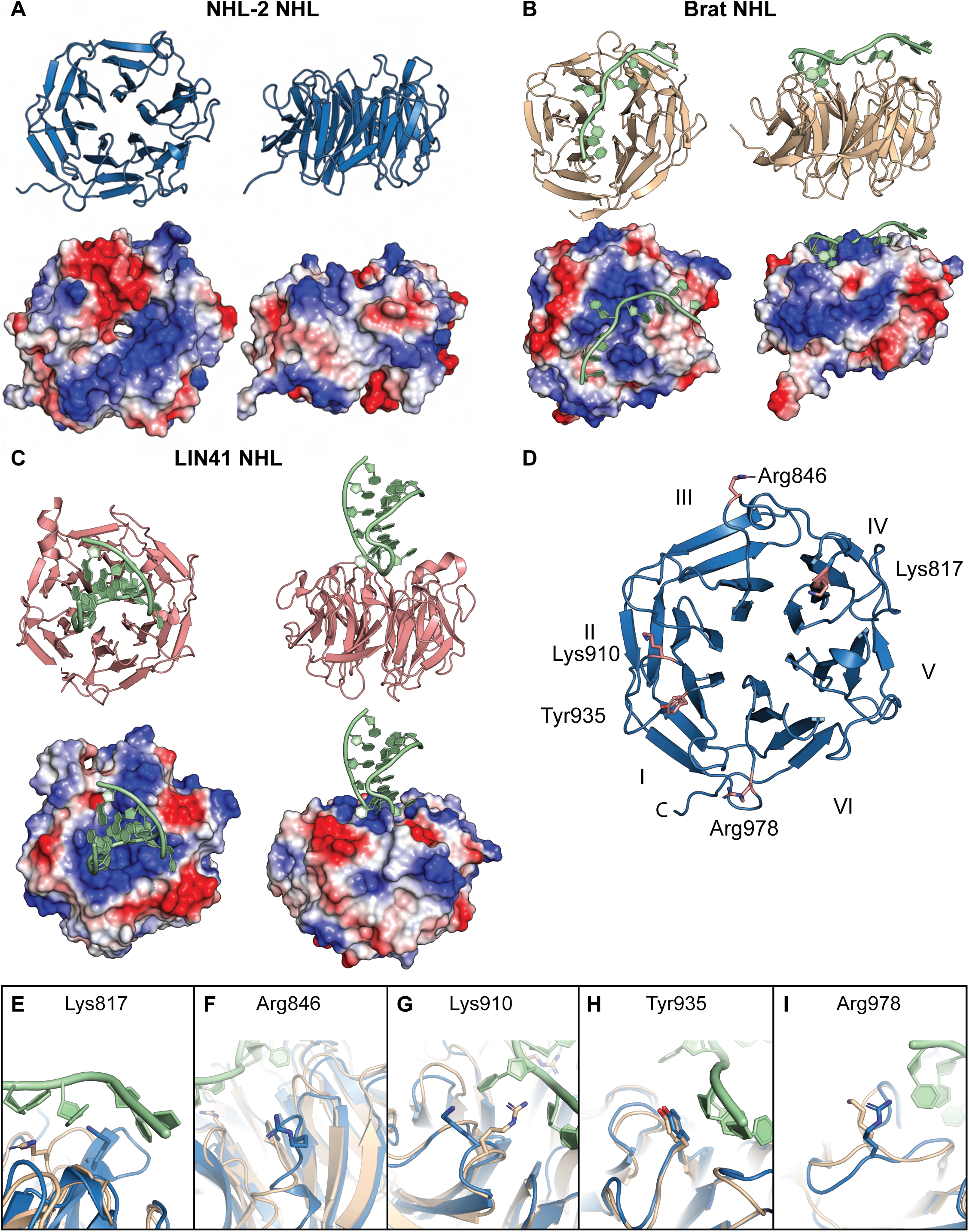
Schematic representation of NHL-2 NHL domain. **A)** The overall NHL domain structure is shown in blue ribbon representation from the “top” of the 6-bladed beta propeller and from the “side” (PDBID:8SUC). Also shown are electrostatic surface potential diagrams from the same orientations. **B)** The dmBrat NHL (PDBID: 4ZLR) and **C)** drLIN41 (PDBID: 6FQ3) structures solved in the presence of RNA are shown alongside in comparable styles. **D)** The overall NHL domain structure is shown in blue ribbon highlighting the 6 blades and the position of 5 amino acids Lys817, Arg846, Lys910, Tyr935, Arg978. **E-I)** Overlay of NHL-2 and Brat NHL domains highlighting amino acids potentially impacting RNA binding. NHL-2 amino acids (blue sticks; PDBIB:8SUC) and dmBrat NHL amino acids (tan sticks) with bound RNA (green sticks) (PDBID: 4ZLR).

**Table 1.**
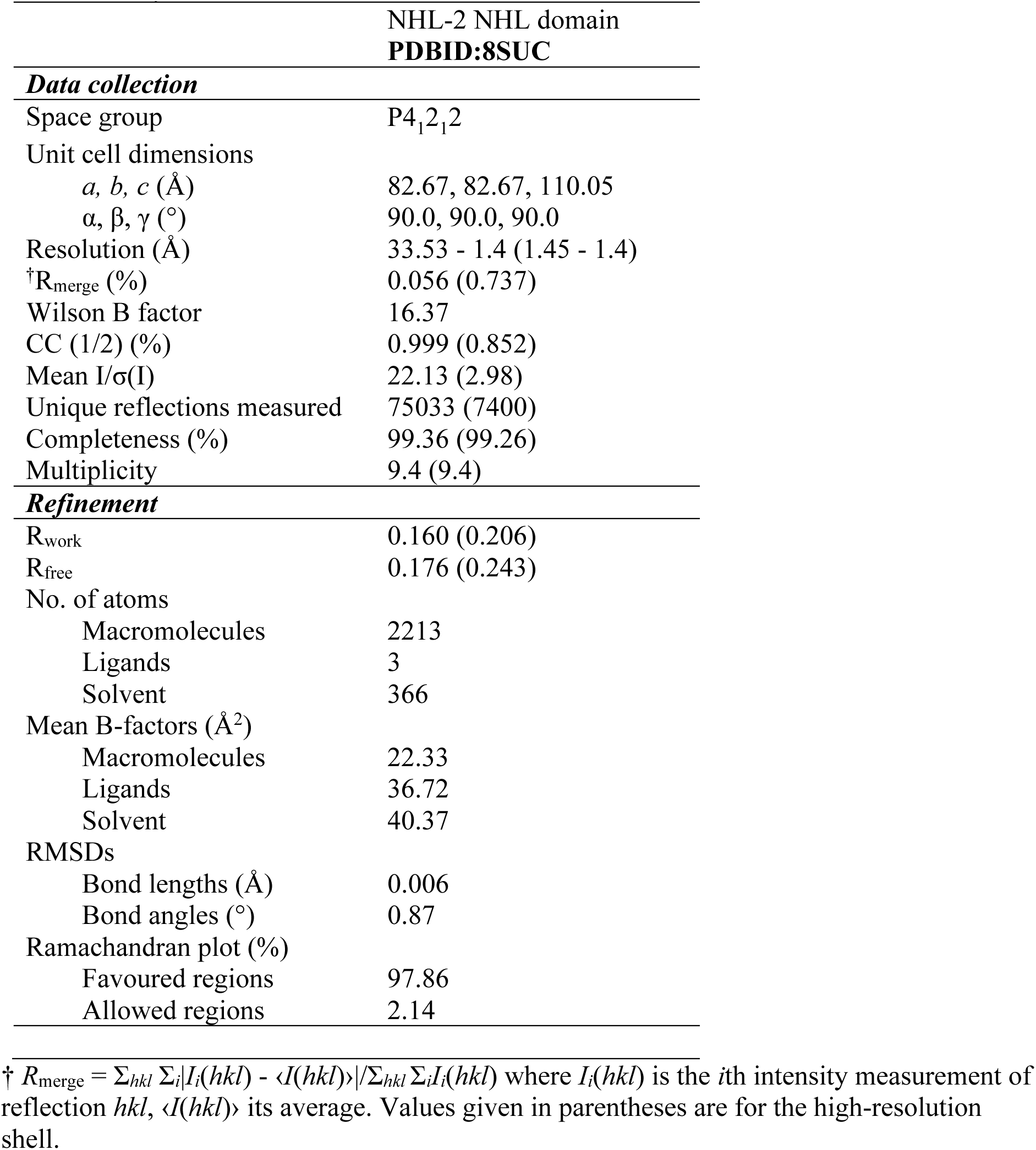
X-ray data statistics.

The top surface of the NHL domain provides a versatile surface for specific recognition and binding of oligonucleotides. This becomes more evident upon inspection of the charged surfaces of the NHL domains. The ceNHL-2 NHL domain (Figure 1A) possesses a striking positively charged groove across blades 6, 1 and 2 which is a likely site for RNA interactions. The adjacent area forms a highly negatively charged patch across blades 3 and 4. In contrast, the dmBRAT and drLIN41 NHL domains possess more centrally located positively charged patches on their “top” surfaces without such a large adjacent negatively charged region. The crystal structures of dmBRAT and drLIN41 NHL were both solved in complex with the target RNA, showing the way in which RNA engages the positively charged regions at the “top” surface of their 6-bladed beta propellers. Interestingly, RNA engages in different modes in these two proteins. The UUGUUG RNA sequence binds in an extended conformation across dmBRAT NHL (25), whereas the GGAGUCCAACUCC sequence that forms an RNA loop structure binds to drLIN41 NHL with a much smaller area of contact with the protein surface (24). Thus, while ceNHL-2 NHL is likely to interact with U-rich RNA at its “top” positively charged patch, the actual orientation and extent of interaction with the protein cannot be reliably predicted.

We were not able to crystallize the ceNHL-2 NHL domain in the presence of 5, 7 or 10mer length U-rich RNA, and therefore sought to identify the RNA-binding site by mutational analysis. Detailed inspection of the ceNHL-2 NHL structure identified 5 amino acids (Lys817, Arg846, Lys910, Tyr935, and Arg978) at the “top” surface of the NHL domain that were potentially in a position to form interactions with bound RNA (Figure 1D). These amino acids were either conserved with dmBRAT residues in key positions engaging RNA or were positively charged amino acids in the vicinity of the positively charged patch of the NHL-2 NHL domain (Figure 1A, D-I).

Therefore, we cloned alanine mutants of the NHL-2 NHL domain at these five amino acid residue positions to identify variants of the NHL domain with altered RNA binding capacity. These mutants would not only help to identify the RNA binding site and indicate the mode of RNA interaction by the NHL-2 NHL domain, but could potentially be used in functional screens of NHL-2 to understand the role of RNA binding in RNA pathways in vivo. We initially designed an NHL-2 NHL construct with all five of the positively charged putative RNA-binding amino acid residues mutated to alanine, referred to as Mut[1-5]. The Mut[1-5] protein was overexpressed and purified as a GST-fusion protein, and its binding to target 17-mer U-rich RNA was measured using fluorescence anisotropy (Figure 2). The wild-type NHL domain bound with low micromolar affinity to the U-rich 17-mer (KD= 0.02 µM ± 0.00). Mut[1-5] showed no ability to bind to RNA at concentrations below 10 µM, representing an effectively abolished RNA binding. An accurate affinity could not be calculated owing to this low binding, but could be estimated to be greater than KD = 62 µM. Thus, these five mutations affected the amino acid residues important for RNA binding.

**Figure 2.**
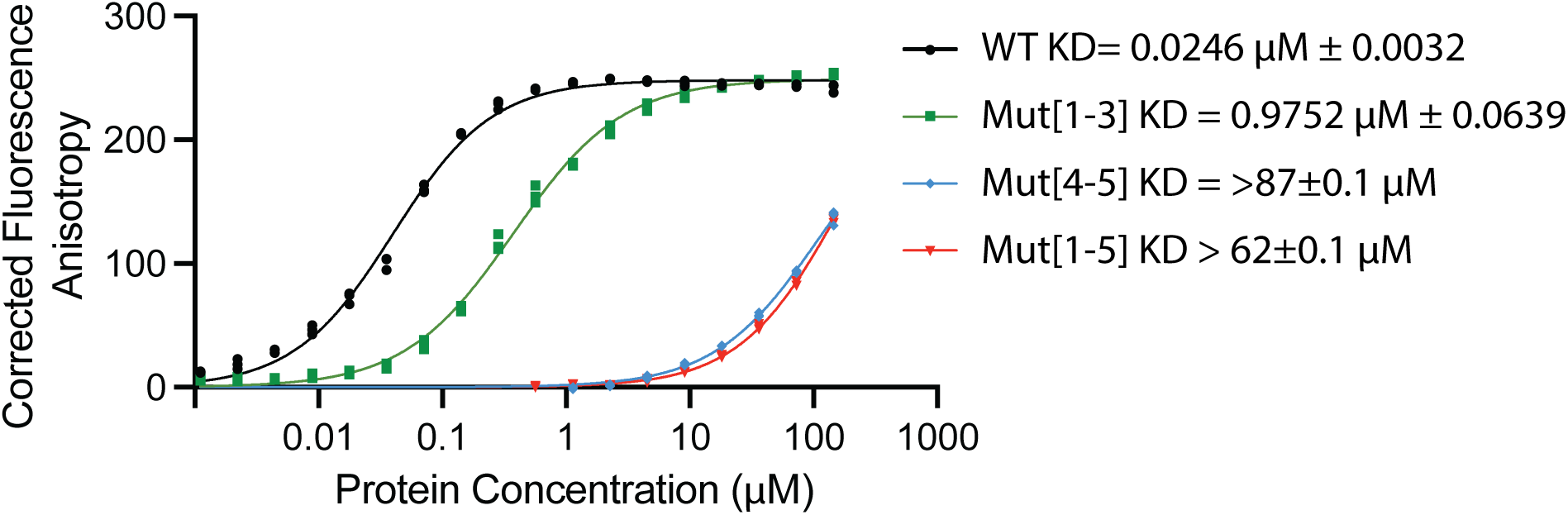
Tyr935 and Arg978 are essential for RNA-binding. Representative fluorescence anisotropy binding curves for wild-type NHL-2 NHL domain and mutant version to 17-mer U-rich RNA. Mut[1-5] (Lys817Ala, Arg846Ala, Lys910Ala, Tyr935Ala, and Arg978Ala), Mut[1-3] (Lys817Ala, Arg846Ala, Lys910Ala) and Mut[4-5] (Tyr935Ala, and Arg978Ala). Data points represent triplicate samples with standard error of the mean (SEM). The binding data were fit by the 1:1 Langmuir binding model to derive the KD.

To further identify the amino acid residues underlying RNA binding, two further mutants were generated: Mut[1-3] involving mutation of Lys817, Arg846, and Lys910 to alanine, and Mut[4-5] involving mutation of Tyr935 and Arg978 to alanine. This division was made on the basis of Mut[1-3] residues being encoded on exon 7 of the gene and Mut[4-5] residues being encoded on exon 8. Again, GST-tagged NHL mutant proteins were generated, and binding to U-rich 17-mer RNA was assessed using fluorescence anisotropy. It was found that Mut[1-3] showed an approximately 50-fold reduced RNA binding ability compared to wild-type NHL, with KD = 0.98 µM ± 0.06. Mut[4-5], on the other hand, showed an abolished RNA binding ability equivalent to that of Mut[1-5] with KD estimated to be greater than KD=87 µM. This reveals that Tyr935 and Arg978 are primarily responsible for NHL-2 NHL binding to RNA, and their mutation to alanine is sufficient to effectively abolish RNA binding.

### RNA-binding is essential for NHL-2 function in the Heterochronic Pathway

To better understand the requirements of the RING and NHL domains for NHL-2 function, we used CRISPR/Cas9 to generate mutants of these domains. We removed the entire RING domain by deleting exons 1 and 2, hereafter referred to as *nhl-2(ΔRING)*. This deletion results in the use of the ATG found at the start of exon 3 as the start codon, and is expected to produce a truncated NHL-2 protein lacking the RING domain but containing all other domains. Expression of the allele was detected using RT-PCR (data not shown). To analyze the requirement for RNA-binding activity of the NHL domain, we mutated Y935 and R978 to alanine, as these mutations abolished RNA binding.

To analyze NHL-2 function, we focused our attention on alae formation as it has been shown to provide a clear readout of NHL-2 function in the *let-7* family regulated heterochronic pathway (26). Alae formation is essential normal in *nhl-2* and *cgh-1* null mutants but is 100% defective in *nhl-2;cgh-1* double null mutant background, suggesting that NHL-2 and CGH-1 are co-dependent for *let-7* miRNA function (26). Therefore, we used *cgh-1(RNAi)* in wild-type, *nhl-2(ok818)* null*, nhl-2(ΔRING)* and *nhl-2(RBlf)* worms to examine the functional requirements of these domains. The *nhl-2(ΔRING)* mutant showed no alae defects by itself; however, *cgh-1*(RNAi) in this strain resulted in 65% of the worms displaying misshapen alae (Figure 3A, B). Similarly, the *nhl-2(RBlf)* mutant had little effect on alae formation (96% normal alae), whereas *cgh-1* RNAi in this background resulted in a major decrease in normal alae formation (15% normal alae) and 45% of worms with no visible alae (Figure 3A, B). This is similar to the effects observed in *nhl-2(ok818)* and *cgh-1(RNAi)* worms (Figure 3A, B). Analogous results were obtained when we tried to rescue the *nhl-2(ok818)* null mutant by generating transgenic worms expressing GFP-NHL-2 with a non-functional RING domain (by replacing Cys77, Cys82, and Cys85 with alanine) or lacking RNA-binding activity (by replacing K817, R846, K910, Y935, and R978 with alanine) (Supplementary Figure 1). Together, these data indicate that the RING and NHL domains are required for the normal function of NHL-2, but that the loss of RNA-binding activity has a more pronounced phenotype, suggesting that RNA-binding is critical for NHL-2 function.

**Figure 3.**
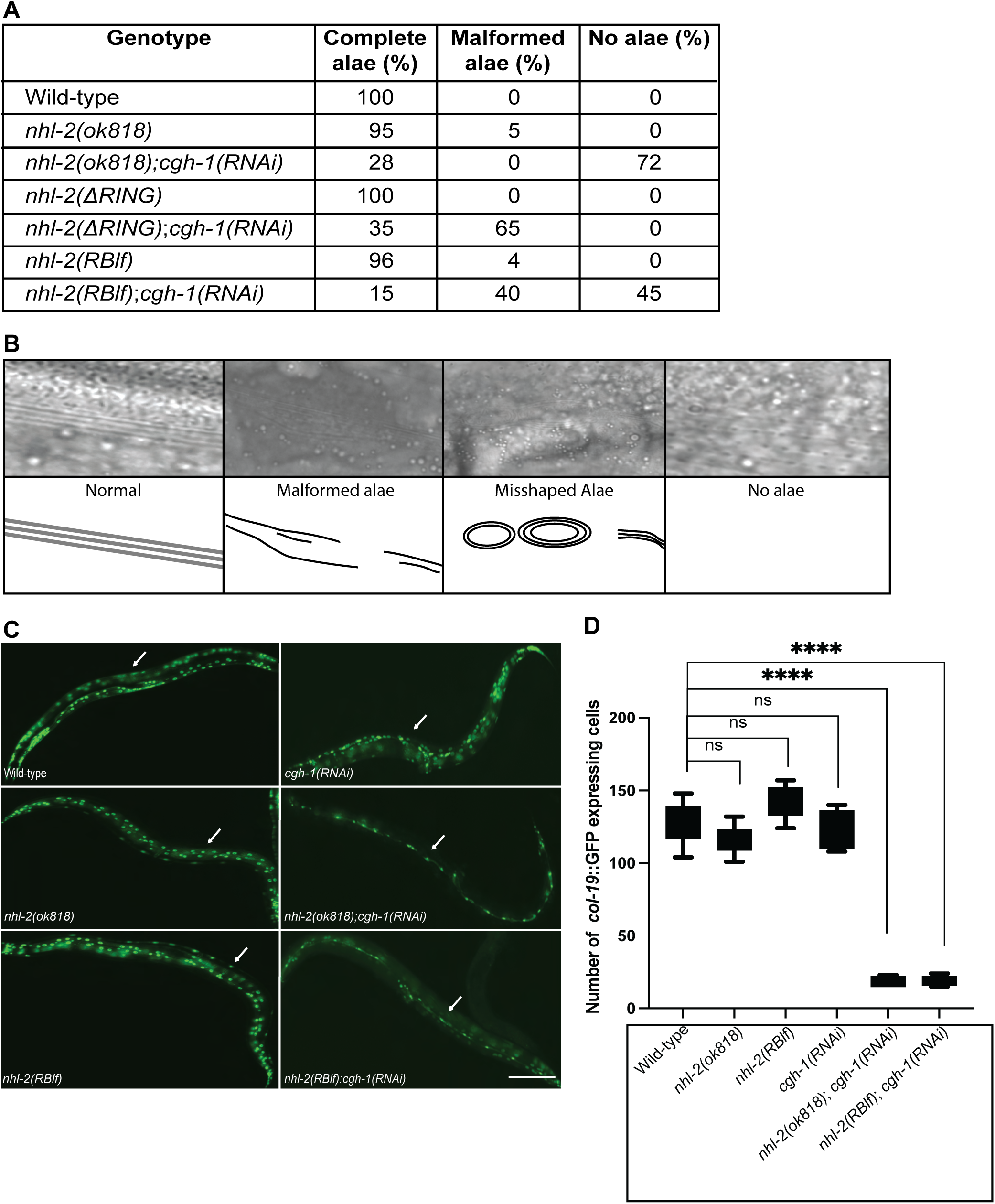
Analysis of genetic interactions of *nhl-2(ΔRING)* and *nhl-2(RBlf)* mutants. **A)** Quantification of alae defects in *nhl-2(ΔRING)* and *nhl-2(RBlf)* worms with and without *cgh-1(RNAi)* **B)** Representative DIC images of normal and defective alae. **C)** Analysis of *col-19*::GFP reporter expression in wild-type, *nhl-2(RBlf)* and *nhl-2(ok818)* worms. **D)** Quantification of *col-19*::GFP reporter expressing cells counted in WT worms and *nhl-2 (ok818), nhl-2(RBlf)* and effect of *cgh-1(RNAi)* on the strains.

We next examined the development of hypodermal cell differentiation using a *col-19*::GFP reporter, as *let-7/*NHL-2 is required for normal expression (26). We generated a strain containing the *col-19*::GFP reporter in the *nhl-2(RBlf)* background and used the *nhl-2(ok818);col*-19::GFP strain (26). As expected, *nhl-2(ok818);col*-19::GFP;*cgh-1(RNAi)* adult worms showed the *col-19*::GFP under-expression phenotype (Figure 3 C, D), which is similar to the triple mutant *nhl-2(ok818):mir*-48;*mir*-84 (26). The *nhl-2(RBlf);col*-19::GFP;*cgh-1(RNAi)* worms exhibited a similar *col-19*::GFP under expression phenotype as *nhl-2(ok818);col*-19::GFP;*cgh-1(RNAi)* (Figure 3C, D), indicating that the RNA-binding activity of NHL-2 is essential for the regulation of the *col-19*::GFP reporter.

To further analyze the requirements of the RING and NHL domains of NHL-2, we examined the synthetic interaction between NHL-2 and other miRISC factors on alae formation (Figure 4). RNAi knockdown of *alg-1* and *ain-1* resulted in a strong alae defect by themselves and was not enhanced in the *nhl-2(ΔRING)* mutant, whereas *nhl-2(RBlf)* and *nhl-2(ok818)* both displayed a significant increase in alae defects. We also knocked down the CGH-1 interacting protein IFET-1 (27), which is an eIF4E binding protein and translational repressor IFET-1 (28). Knockdown of *ifet-* 1 in wild-type worms resulted in over 50% of worms with alae defects, similar to the knockdown of *alg-1* and *ain-1*. This defect was not increased in *nhl-2(ΔRING)* mutant, while *nhl-2(RBlf)* and *nhl-2(ok818)* showed significantly increased alae defects, suggesting that IFET-1 is a critical component of miRISC effector complex. EDC-3 is a conserved decapping enhancer factor that interacts with CGH-1 in a yeast two-hybrid system (29). A small but significant synthetic alae defect was observed when *edc-3* was knocked down in the *nhl-2(ΔRING)* mutant, and this defect was further enhanced in the *nhl-2(RBlf)* and *nhl-2(ok818)* backgrounds. The *C. elegans* casein kinase II (CK2) homologue KIN-3 aids miRISC binding to mRNA targets and activates CGH-1 by phosphorylating it (Alessi et al., 2015). Knockdown of *kin-3* in the *nhl-2(RING)*, *nhl-2(RBlf)* and *nhl-2(ok818)* backgrounds all showed alae defects. NTL-3 is part of the CCR4-NOT transcriptional regulation complex in *C. elegans,* is a yeast paralog of Not3p and Not5p (30) and was identified in a synthetic RNAi screen for NHL-2 interactors that cause fertility defects (31). Knockdown of *ntl-3* resulted in alae defects in wild-type worms, which was not enhanced in the *nhl-2(ΔRING)* mutant, whereas the *nhl-2(RBlf)* and *nhl-2(ok818)* backgrounds both showed significant increases in alae defects. TEG-1 regulates the stability of VIG-1 and ALG-1 containing miRISC and is required for *let-7*-mediated regulation (32). Knockdown of *teg-1* resulted in an alae defect that was not enhanced in the *nhl-2(ΔRING)* mutant, whereas *nhl-2(RBlf)* and *nhl-2(ok818)* both displayed a significant increase in alae defects. VIG-1 is a conserved protein that has been biochemically detected as part of the miRISC subcomplexes and is associated with *let-7* microRNA-binding complexes in *C. elegans* (33). Knockdown of *vig-1* induced a relatively minor alae defect only in the *nhl-2(RBlf)* and *nhl-2(ok818)* backgrounds. Together, these experiments support the conclusion that *nhl-2(RBlf)* and *nhl-2(ok818)* have synthetic interactions with some, but not all, miRISC factors and that RNA binding is essential for its function.

**Figure 4.**
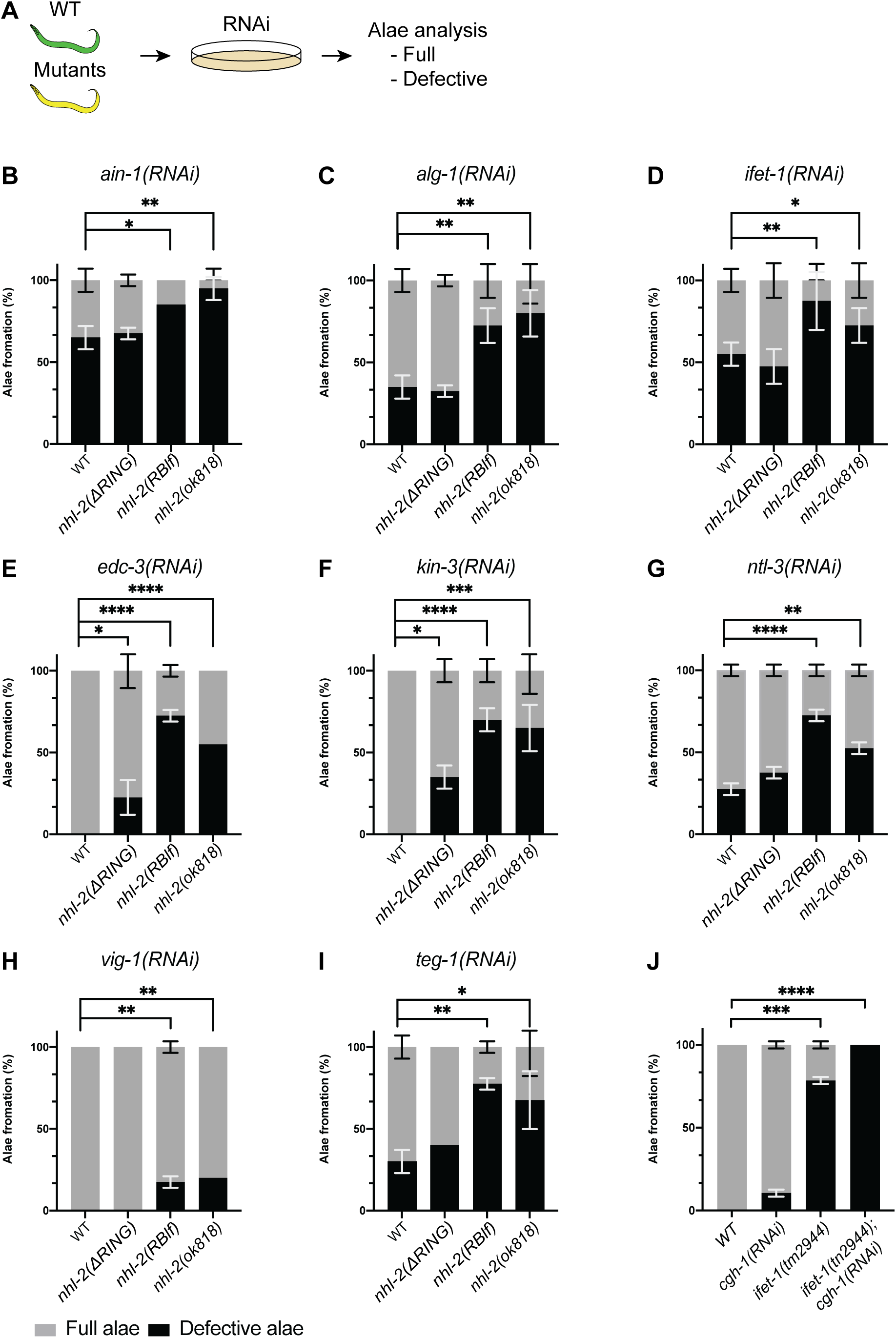
Synthetic interaction of *nhl-2* mutants with selected miRISC cofactors. **A)** Schematic diagram of experimental design. **B)** Quantification of alae formation in wild-type, *nhl-2(ΔRING)* and *nhl-2(RBlf)*, *nhl-2(ok818)* worms were determined when specific miRISC cofactors were knocked down. Alae were scored normal or defective (combined mis-shaped and no alae). Error bars represent the standard deviation.

Given the strong interaction between *ifet-1* and *nhl-2*, we next analyzed alae formation in the *ifet-1(tm2944)* null mutant with and without knockdown of *cgh-1.* The *ifet-1* null worms had a high penetrance of defective alae (gapped 76%), which was significantly enhanced when *cgh-1* was knocked down (100% defective alae, of which 88% of worms had no alae) (Figure 4I). Together, these data suggest that IFET-1 is a component of the miRISC effector complex that is required for alae formation.

### Putative NHL-2 binding sites are required for *lin-28* reporter gene regulation

Having established that RNA binding is critical for NHL-2 function, we next investigated whether the putative NHL-2 binding U-rich motif (UUUUUU) (22) is required for the regulation of *let-7* family miRNA targets. We examined the 3’ UTRs of several well-characterized targets, *daf-12*, *hbl-1*, *let-60* and *lin-28*, and putative NHL-2 binding sites were identified in all 3’ UTRs. However, there was no obvious pattern of their location in the 3’ UTR or proximity to the *let-7* family binding site (data not shown). We focused our attention on *lin-28* as it has the shortest 3’ UTR and contains the fewest NHL-2 binding putative sites compared to other *let-7* family targets.

To examine the requirement of putative NHL-2 RNA-binding motifs in *lin-28* 3’ UTR, we designed a seam cell-specific reporter construct (Figure 5A). This reporter uses the *col-10* promoter *(pcol-10)* to drive seam cell-specific expression of the mCherry::H2B protein. To ensure that any observed mCherry::H2B is the result of recent translation and not carryover of the earlier translated protein, we tagged mCherry::H2B with the PEST domain, which significantly reduce protein half-life (34). Given that *lin-28* is regulated by both *let-7* family and *lin-4* miRNAs (35,36), we mutated three nucleotides in the *lin-4* seed sequences to remove the inhibition via this pathway, leaving the mCherry::H2B reporter only under the control of the *let-7* family miRNA (36). We developed two variations of the base construct. One contained the putative NHL-2 binding sites and the second included single or double cytosines in the middle of the U-rich motif, as this was not bound by the RNA-binding domain of NHL-2 (Boag/Wilce labs data not shown) (Figure 5A). As expected, fluorescence microscopy of transgenic worms revealed that both constructs were expressed in the L1 stage worms in both wild-type and *nhl-2(ok818)* worms, as *let-7* miRNA did not affect *lin-28* at the L1 stage (36). Wild-type worms carrying transgenes containing the wild-type construct showed no observable mCherry at the adult stage as expected. In contrast, transgenic wild-type worms with the mutated putative NHL-2 binding sites displayed mCherry expression at the adult stage (Figure 5). Interestingly, adult *nhl-2(ok818)* worms containing the wild-type and mutated putative NHL-2 binding sites construct expressed mCherry. Together, these data demonstrated that NHL-2 and the putative NHL-2 RNA-binding sites in the 3’ UTR of *lin-28* are important for normal regulation translation.

**Figure 5.**
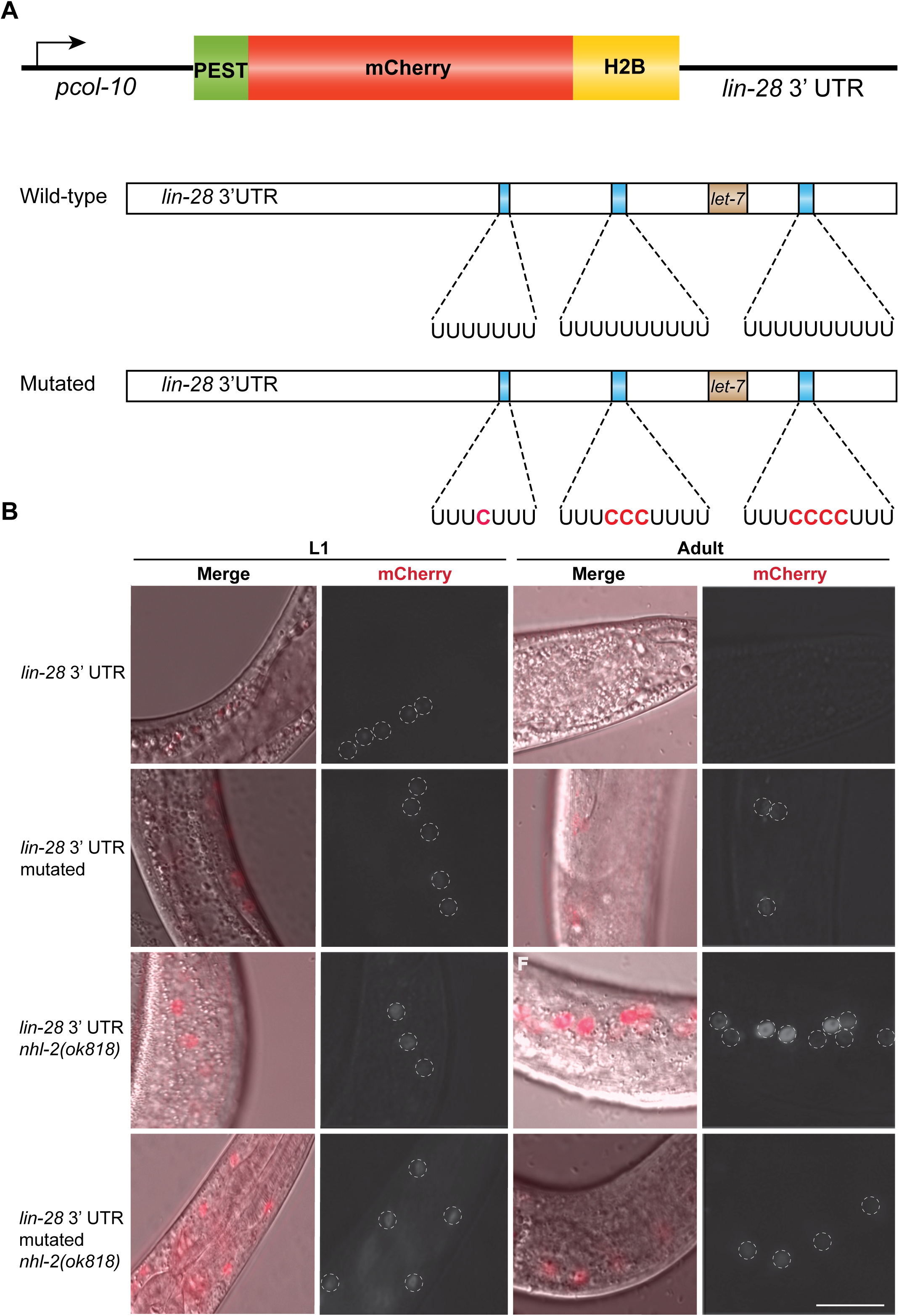
Putative NHL-2 binding sites are required for *lin-28* 3’ UTR Reporter regulation. **A)** Schematic representation of the PEST-mCherry-H2B reporter constructs. The p*col-10* promoter drives expression of the PEST::mCherry-H2B reporter in seam cells and translation regulated via the *lin-28* 3’ UTR. The *let-7* binding site is shown in brown. The wild-type *lin-28* 3’ UTR contains three putative NHL-2 binding sites which were mutagenized by addition of the cytosine into the middle of the binding site. **B)** Analysis of *lin-28* 3’ UTR mCherry reporter constructs in seam cells of wild-type and *nhl-2(ok818)* worms.

### P-body size is increased in NHL-2(RBlf) worms

An unexpected observation in GFP::NHL-2(RBlf) expressing worms was the appearance of large cytoplasmic foci in some somatic cells resembling P-bodies. To confirm that GFP::NHL-2(RBlf) worms had increased P-body size we generated a strain that expressed both GFP::NHL-2(RBlf) and mRuby::DCAP-1 (37) as a P-body marker. We analyzed an average of 45 P bodies in the tail of 15 one-day-old adult worms for each strain using fluorescence microscopy and found a significant increase in the size of the P bodies in NHL-2(RBlf) compared to wild-type (2 µm compared to 1 µm (*P* value <0.0001) (Figure 6A, B). Similar results were obtained when we expressed *nhl-2-2(RBlf)* extrachromosomal arrays in the *nhl-2(ok818)* background (data not shown) Interestingly, extrachromosomal array expression of the mutated RING domain NHL-2 was still localized to P-bodies; however, P-body size did not increase (data not shown). P-body size has been shown to vary when mRNA decay pathways are altered (38) therefore, a possible reason for the increase in the size of P-bodies in NHL-2(RBlf)-expressing worms is a defect in miRNA-mediated turnover of target mRNAs. To further examine P-body dynamics in wild-type and NHL-2(RBlf) worms, we used RNAi to knock down key factors in deadenylation and decapping-mediated mRNA decay pathways, as they have been shown to influence P-body size (38). We targeted the deadenylating factors CCR4, CCF-1, and PAN2, the decapping activator factors PATR1 and CGH-1, and exoribonuclease XRN-1. As expected, RNAi knockdown of the deadenylation factors *panl-2, ccr4, and ccf-1* in GFP::NHL-2 worms resulted in decreased P-body size. This is consistent with previous work showing that defects in deadenylation result in reduced mRNA turnover and P-body size (39,40). No significant change in P-body size was observed in *cgh-1* and *patr-1* RNAi wild-type worms (Figure 6C), whereas *xrn-1* RNAi caused an increase in the size of P-bodies in wild-type worms (Figure 6C). Consistent with the GFP::NHL-2 results, the size of P-bodies was reduced in RNAi-knocked down *panl-2* and *ccf-1* cells in the GFP::NHL-2(RBlf) background (Figure 6).

**Figure 6.**
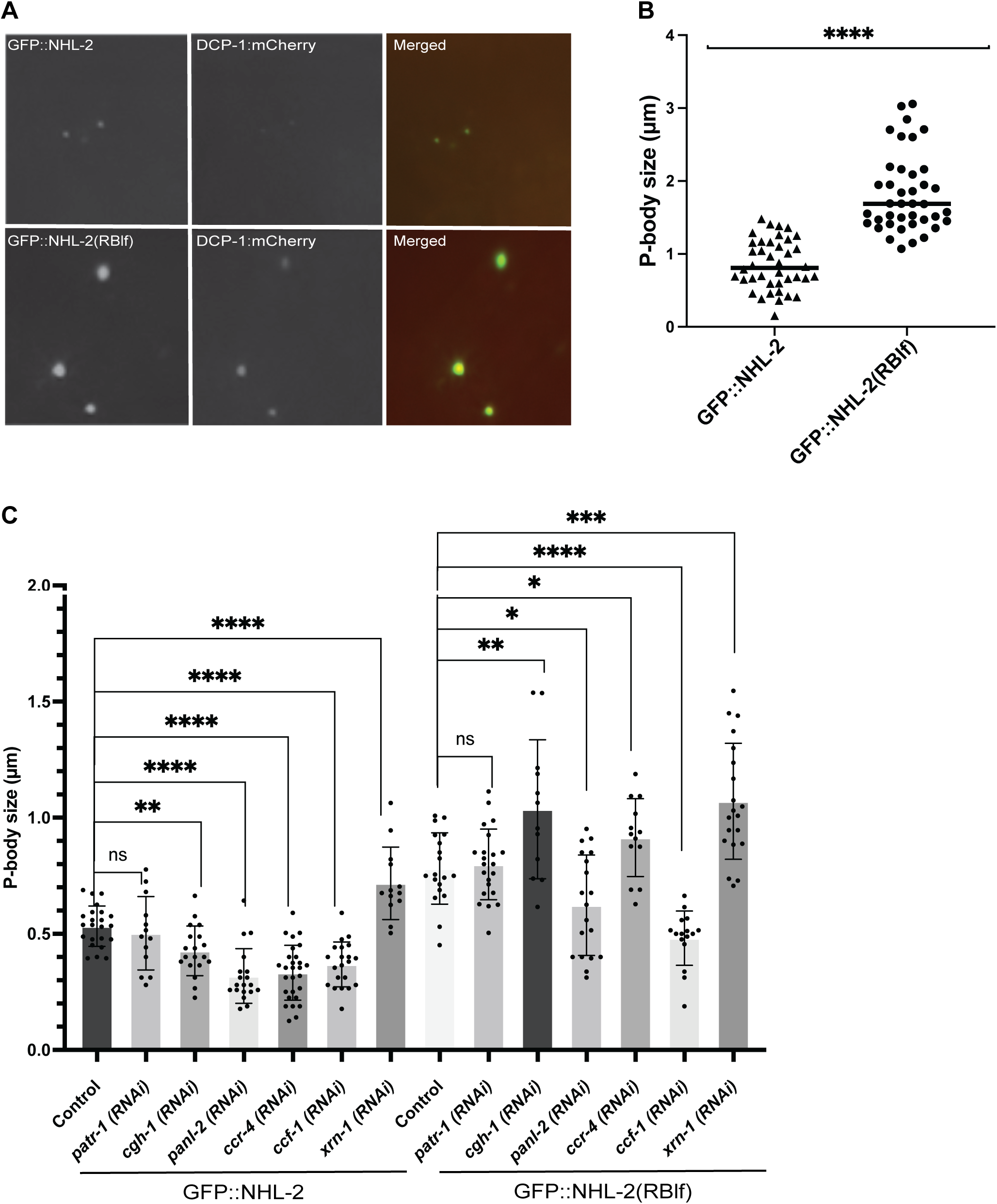
P bodies are larger in NHL-2(RBlf) worms. **A)** GFP::NHL-2 and GFP::NHL-2(RBlf) co-localise with mCherry::DCAP-1 in P-bodies, however GFP::NHL-2(RBlf) P-bodies are larger compared to GFP:: NHL-2. **B)** Quantification of P-bodies in GFP::NHL-2 and GFP::NHL-2(RBlf). P-body sizes were measured in 15 adult worms in the tail area. *P* value <0.0001. **C)** Quantification of P-body size in L4 worms expressing GFP::NHL-2 and NHL-2(= RBlf) subjected to RNAi knockdown of specific RNA decay factors. Images were analysed using Fiji software to determine P-body size. Data were analysed using one-way ANOVA multiple by GraphPad Prism 9.5. P-bodies were measured in 5 worms in each strain.

Interestingly, the knockdown of *ccr*4 in NHL-2 RNA-binding mutant worms resulted in a small but significant increase in P-body size, which was different from the wild-type NHL-2 worms. There was no significant change in *patr-1* RNAi results when comparing the size of P-bodies to control RNAi, which phenocopied GFP::NHL-2 worms. The size of P-bodies in *xrn-1*(RNAi) in *nhl-2(RBlf)* increased, consistent with what was observed in GFP::NHL-2 worms (Figure5C). Interestingly, P-body size increased in *cgh-1(RNAi)* in *nhl-2(RBlf)* mutant compared with GFP::NHL-2 worms (Figure 6C), which is consistent with the strong synthetic interactions observed with NHL-2 and CGH-1. Collectively, these data indicate that although the size of P-bodies in NHL-2(RBlf) are larger than normal, they are still influenced by the action of deadenylation factors and decay factors, similar to wild-type worms. This suggests that the increased size of NHL-1(RBlf) P-bodies is likely to reflect alterations in the rate of mRNA decay.

### Increased association of *let-7* family target mRNAs with ALG-1/-2 in nhl-2(ok818) worms

Next, we wanted to understand if the association of *let-7* target mRNAs with miRISC is altered in the absence of NHL-2. To do this, we crossed *nhl-2(ok818)* and worms with a strain containing FLAG-tagged ALG-1 and ALG-2 (41). Given the clear genetic interaction between *nhl-*2 and *cgh-* 1, we knocked down *cgh-1* by RNAi in *nhl-2(ok818);*FLAG::ALG-1/2 and wild-type (FLAG::ALG-1/2) worms to determine whether this altered *let-7* target mRNAs associated with ALG-1/-2. ALG-1 and ALG-2 were immunoprecipitated using an anti-FLAG antibody, and co-immunoprecipitated RNAs were extracted and quantified by qRT-PCR. RNA was also extracted from the protein lysate prior to IP to determine total mRNA levels. We chose *daf-12*, *hbl-1*, *let-*60, and *lin-41* to characterize as they represent some of the best described *let-7* targets. Strikingly, in all cases, there was at least a two-fold increase in the mRNAs in the ALG-1/-2 IPs for all four mRNAs quantified (Figure 7). Interestingly, *cgh-1(RNAi)* in the wild-type background did not affect the association between *let-7* target mRNA and ALG-1/-2. Similarly, no change in enrichments was observed in the *nhl-2(ok818)* background with or without *cgh-1(RNAi)*. When we looked at the total mRNA level, they were not significantly affected in *nhl-2(ok818)* or when *cgh-*1 was knocked down in wild-type or *nhl-2(ok818)* worms. Together, these data suggest that an increased number of AGO-bound mRNAs is not due to an increase in total mRNA levels, but the result of more mRNAs being bound by AGOs. A likely explanation for this is that AGO-1/-2 bound mRNAs are not degraded as efficiently by the deadenylation and decapping machinery in the absence of NHL-2.

**Figure 7.**
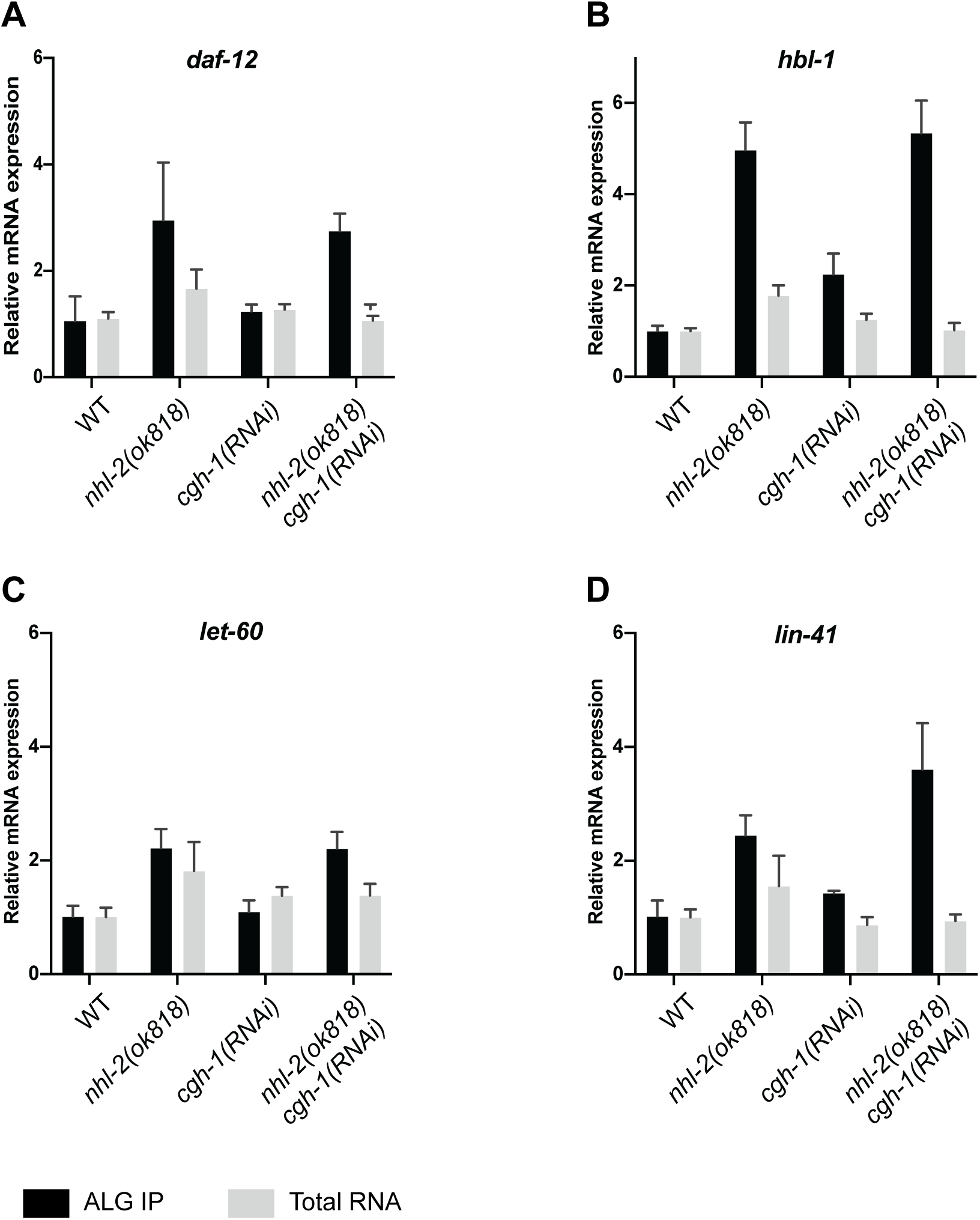
*let-7* target mRNAs are enriched in ALG1/2 IPs. Fold of enrichment of *let-7* target mRNAs *daf-12, hbl-1, let-60 and lin-41* in the ALG1/2 IPs and total RNA from L4 stage wild-type (WT), *nhl-2(ok818), cgh-1(RNAi)* and *nhl-2(0ok818);cgh-1(RNAi)* worms. For total RNA samples, mRNA levels were quantified by qRT-PCR and normalized to *act-1* mRNA. For RNA isolated from the ALG-1/ALG-2 IPs, fold enrichment of a given mRNA was determined by comparing the normalized concentration of this mRNA in the IP sample to the corresponding input worm lysate sample. Error bars represent the 95% confidence intervals calculated from samples using the ΔΔCt method.

## Discussion

In this study, we determined the crystal structure of the RNA-binding domain of NHL-2 and identified key amino acids for RNA-binding and functional testing of the requirement of RNA-binding activity for NHL-2 function. Multiple lines of evidence suggest that the absence of functional NHL-2 results in changes in RNA fate, including increased P-body size and an increased association of some miRNA target mRNAs with the miRISC. Overall, our data are consistent with a model in which NHL-2 functions in conjunction with CGH-1, IFET-1, and its directly interacting protein IFE-3 (eIF4E) to inhibit the translation and mRNA turnover of some miRNA targets.

### NHL-2 Crystal structure reveal similarities and differences to other NHL domains

The NHL domain is an evolutionarily conserved fold in a β-propeller architecture that can function as an RNA-binding (21,24,25) and/or protein-protein interaction platform (42). The structure of the NHL-2 NHL domain provides another example of an NHL domain featuring a large positively charged patch on the “top” surface that facilitates RNA-binding (24,25). In contrast to BRAT NHL and Lin41 NHL domains, the positively charged patch is strikingly elongated and extends to the 6^th^ blade of the propeller. The mode of RNA binding by NHL-2 NHL is likely to be more similar to that of BRAT NHL bound to an elongated stretch of RNA than that of LIN41 NHL bound to an RNA loop structure. Indeed, NHL-2 NHL amino acids Y935 and R978, identified as essential to RNA binding and situated across blades 5 and 6, correspond to amino acids Y959 and K998 in BRAT NHL that may be similarly involved in RNA binding. Future structural studies of NHL-2 NHL in complex with RNA will hopefully reveal the binding mode and basis of specificity of NHL-2 NHL for detailed comparison with BRAT, LIN41 and other NHL structures solved in complex with RNA (24,25,43).

The RING domains are often associated with E3 ubiquitin ligase activity (44,45). The RING domain of TRIM32 is required for the regulation of mouse neural progenitor cells via ubiquitination of the transcription factor c-Myc but is not required for miRNA-mediated repression (18). Similarly, the RING domains of LIN-41 in *C. elegans* (46) and human TRIM71 (47) are not required for their function, indicating that there is a strong cellular context for E3 ubiquitin ligase activity. Although the *nhl-2(ΔRING)* mutant displayed some developmental defects, it was significantly milder than *nhl-2(RBlf)*, suggesting that this domain is not essential for NHL-2 function. One caveat to this conclusion is that we have not been able to determine the amount of NHL-2 protein in *nhl-2(ΔRING)* mutant worms as the anti-NHL-2 antibody we have is to the deleted N-terminal region of the protein and cannot detect this protein isoform. It is possible that there is reduced NHL-2(*Δ*RING) protein in the worms due to the deletion of exons one and two, which could result in the observed phenotypes. The U-enriched NHL-2 binding motif was found multiple times in the 3’-UTRs of some *let-7* family target mRNAs (data not shown), which could allow multiple NHL-2 proteins to bind. TRIM-NHL proteins have been found to dimerize and tetramerize through the RING (48) and coil-coiled domains (49) which could facilitate the formation of higher-order structures that allow multiple NHL domains to bind to mRNA targets and enhance their substrate affinity (50). It is conceivable that the RING domain of NHL-2 is involved in the formation of dimer/tetramers of NHL-2, which enhances substrate affinity. In the absence of this domain, the reduction in affinity has a modest impact on miRISC activity, which is associated with relatively minor developmental phenotypes.

The increase in the P-body size in NHL-2(RBlf) worms was unexpected. P-bodies are complex heterogeneous structures that were originally thought to be sites of mRNA decay as they are enriched for many decay factors (51). However, more recent studies have found that decay occurs in the cytosol, not in P-bodies (52) and that mRNAs can exit P-bodies and return to translation (53,54). The current model of P-body function suggests that P-bodies are sites where translationally competent mRNAs and their regulators are stored temporarily. Despite the evidence suggesting that mRNA decay occurs outside of P-bodies, P-body size is clearly dependent upon the normal function of the 5’ to 3’ mRNA decay pathways. For example, P-body size increases when the function of 5’ to 3’ exonuclease XRN-1 is inhibited in yeast, *drosophila* and human cells (55–58). Conversely, inhibition of deadenylation results in smaller P-bodies (59). Interestingly, disrupting the EDC4-XRN1 interaction inhibits mRNA decapping, stabilizes miRNA-targeted mRNAs, and increases P-body size in HeLa cells (60). The increase in P-body size in *nhl-2(RBlf)* but not in the *RING* mutant, correlates with the severity of the alae defects observed. This suggests the *nhl-2(RBlf)* mutation may disrupt normal miRNA-mediated mRNA decay, although the precise mechanism remains unclear.

The identification of IFET-1 as a miRISC co-factor links miRISC-mediated mRNA regulation with protein interactions at the 5’ end of mRNAs. *C. elegans* has five eIF4E homologues: IFE-1, IFE-2, IFE-3, IFE-4, and IFE-5, which differ in their 5’ cap-binding specificities (61). A recent study found that the conserved glycine-tyrosine-phenylalanine domain-containing protein GYF-1 specifically interacts with IFE-4 and is a miRISC effector protein (10). We have reported that IFET-1 (4E-T homologue) specifically interacts with IFE-3 (62) and proteomic analysis of ALG-1/ALG-2 IPs found that IFET-1 was significantly enriched (Supplemental Figure 2), supporting its direct role in the translational regulation of some miRNA targets. We identified a strong synthetic interaction between *cgh-1* and *ifet-1* which further supported the NHL-2, CGH-1, and IFET-1 axis. The interaction of the DDX6 family of DEAD box RNA helicases and 4E-Ts is conserved (63,64), and using AlphaFold2 we modelled the interaction between CGH-1 and IFET-1. The AlphaFold2 model suggests that the highly conserved amino acids in the Cup Homology Domain (CHD) of human 4E-T that interact with DDX6_C_ (64) would also support the interaction between IFET-1 and CGH-1 (Figure 8). Together, these data support the conclusion that miRNA activity is not only regulated by the binding of the miRNA to its target mRNA via the seed sequence but can also be strongly influenced by the assembly of miRNA-specific effector complexes, at least in part, by the interaction of the 5’-cap-binding protein and its cognate 4E-T translation repressor. This dual mechanism of functional miRNA effector complex formation provides an extra layer of complexity for miRNA-mediated gene regulation. Many co-factors of the core miRISC have been identified, and some appear to function in specific contexts (10,11,65). It is becoming clear that there are several forms of mature miRISC which contain the core proteins Ago and GW182 and are supplemented by various co-factors that form distinct biochemical complexes. For example, GYF-1 is required for *let-7* and *miR-35,* but is dispensable for *lin-4* and *lsy-6* miRNA function (10), and NHL-2, which is required for *let-7* family and *lsy-6* (this work (26). The impact of specific co-factors on the function of mature miRISCs suggests that there may be a strong cell and developmental function of the miRNA-Ago complex (10,12,66). Phosphorylation of Argonautes and other miRISC components is a major mechanism regulating miRISC activity; for example, phosphorylation by Casein Kinase II (CK2) regulates ALG-1 and CGH-1 activities (67,68). NHL-2, IFET-1, and IFE-3 are extensively phosphorylated (69,70) and could play an important role in coordinating their functions, and exploring this in the context of miRISC function is important.

**Figure 8.**
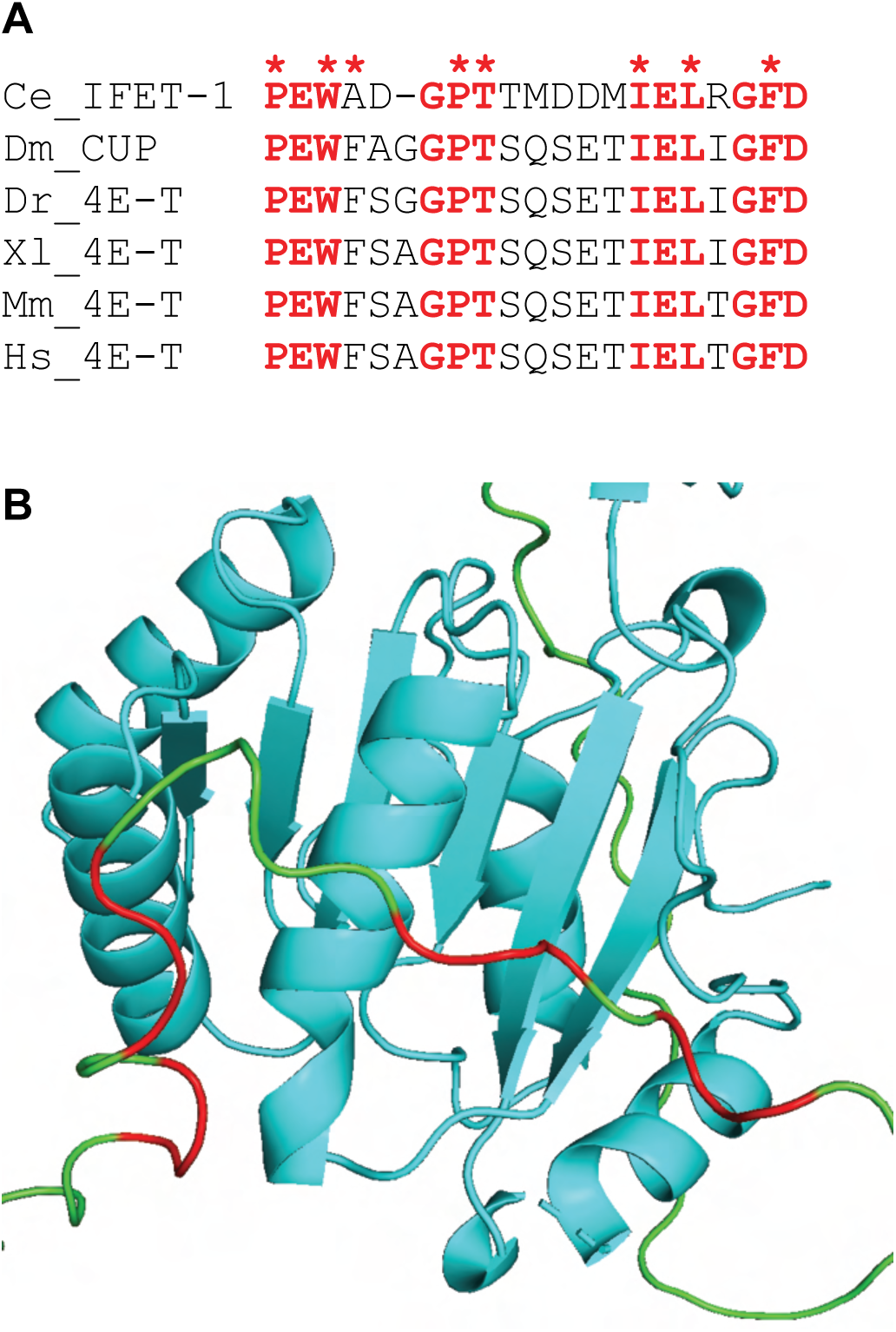
IFET-1 and CGH-1 are predicted to physically interact. **A)** Alignment of the Cup Homology Domain (CHD) from *C. elegans* (Ce), *D. melanogaster* (Dm), *D. rerio* (Dr), *X. laevis* (Xl), M. (Mm) and *H. sapiens* (Hs). Conserved residues are highlighted in red and human 4E-T amino acids that interact with DDX6_C_ are indicated with red asterisks. **B)** AlphaFold2 structure prediction of the IFET-1-CGH-1 interaction. The CGH-1 Rec2A (amino acids 246-420) is shown in aqua, and the CHD of IFET-1 is shown in green with predicted CGH-1 interacting amino acids highlighted in red.

### A model of NHL-2 function

We have shown that NHL-2, CGH-1, and IFET-1 are required for the normal functioning of *let-7* family miRNAs to coordinate normal development. We propose a model in which NHL-2, CGH-1, and IFET-1 work together to translationally inhibit some miRNA targets (Figure 9). In wild-type worms, these proteins work together to inhibit translation and may also help recruit decay factors such as PATR-1 and EDC-3, which competitively bind to CGH-1 (71), the deadenylation machinery PAN-2/PAN-3, and the CCR4-NOT complex. Individual loss of NHL-2, CGH-1, or IFET-1 have a minor effect on miRISC function, while the loss of function of two components significantly impacts miRISC activity, resulting in continued translation of the target mRNA, leading to developmental defects. The role of RNA-binding of NHL-2 remains unclear, but it could be involved in stabilizing the interaction of the NHL-2, CGH-1, and IFET-1 networks on the mRNA, which may help recruit other effector subunits, such as the deadenylation and decay complexes.

**Figure 9.**
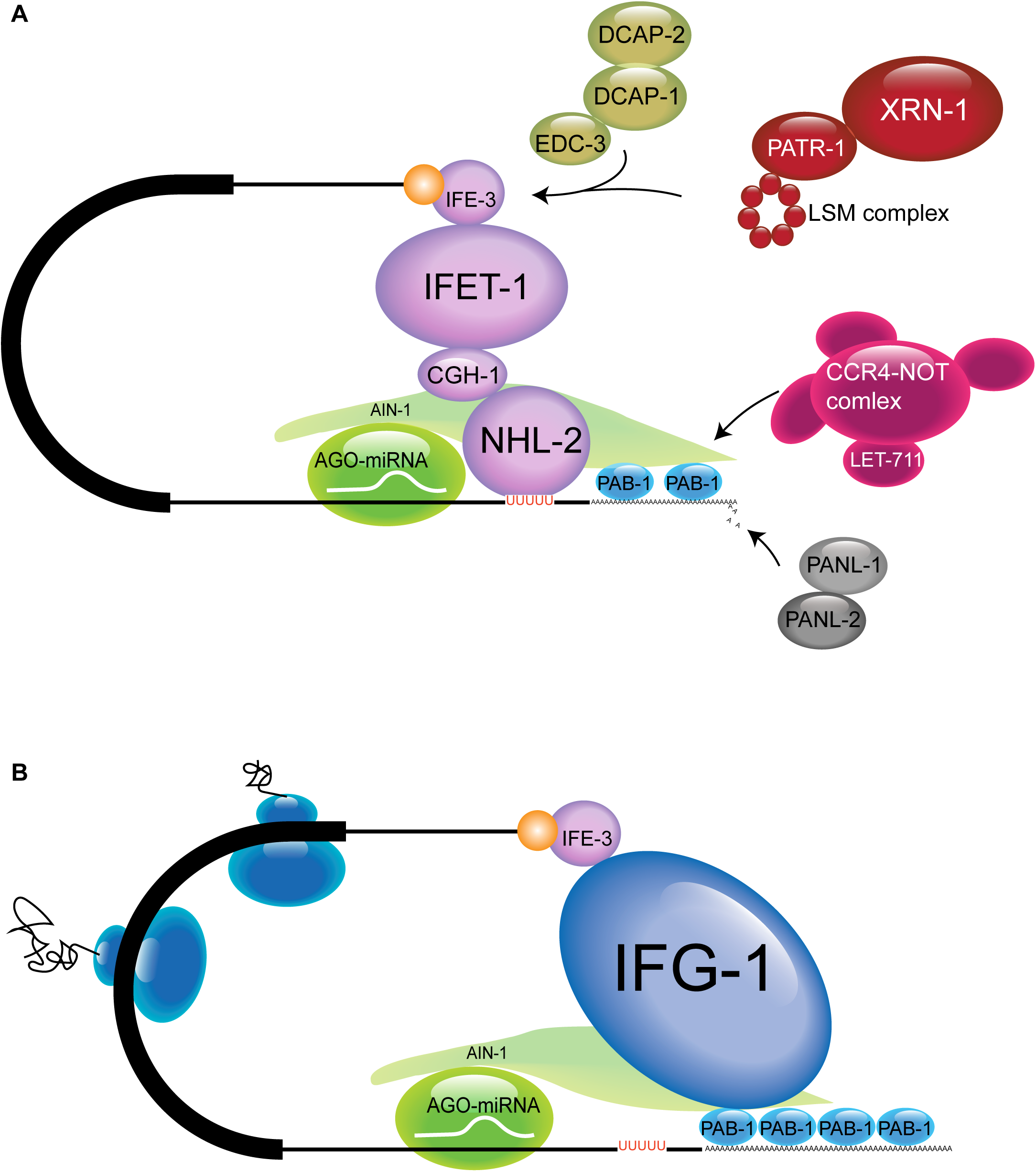
A model of function of NHL-2/CGH-1/IFET-1 effector complex in miRNA-mediated silencing. **A)** In wild-type worms NHL-2 binds to selected miRISC target mRNAs and directly interacts with CGH-1, which in turn directly binds to IFET-1, resulting in inhibition of translation initiation. The direct interaction between IFET-1 and IFE-3 provides a mechanism for selecting subsets of miRNA targets 5′ cap structure. Conserved protein-protein interactions with the NHL-2/CGH-1/IFET-1 effector complex results in the recruitment of the deadenylation and decapping complexes which then mRNA decay. **B)** In the absence of two of the three members of the NHL-2/CGH-1/IFET-1 translational inhibition effector complex the miRNA target remains available for translation.

This work provides a new understanding of the function of some TRIM-NHL proteins in miRNA-mediated mRNA regulation and highlights the key role of NHL-2s RNA-binding. The conserved nature of the NHL-2, CGH-1, and IFET-1 interactions suggests that this could be a mechanism employed in other species. It provides additional evidence that the selection of miRNA targets for transitional repression can be influenced by the mRNA cap-binding protein and its cognate binding partner (IFE-3-IFET-1 and IFE-4-GYF-1).

## Materials and Methods

### *C. elegans* maintenance and genetics

The following strains were used in this study: N2 Bristol (WT), PRB172 (*ifet-1(tn2944)/qC1 [dpy-19(e1259) glp-1(q339) qIs26] III*), PRB310 (*nhl-2(ok818) III*), PRB437 (*hjSi397[dpy-30p::mRuby::dcap-1] I*), (ma371 cdb115 cdb120 [*3xflag::gfp::nhl-2] III)*), PRB438 (*hjSi397[dpy-30p::mRuby::dcap-1] I*), ((ma371 [*3xflag::gfp::nhl-2] III)*), PRB432 (*alg-2(ap431[3xflag::mkate2::alg-2]) II;* (*nhl-2(ok818) III*)*, alg-1(ap423[3xflag::gfp::alg-1]) X*), PRB433 (*qfEX024*(p*col-10*::PEST-mCherry-H2B::*lin-28* 3’ UTR, *rol-6(su1006)*), PRB434 (*qfEX025*(p*col-10*::PEST-mCherry-H2B::*lin-28(mutated)* 3’ UTR, *rol-6(su1006)*), JMC181 (*nhl-2(tor110) III*), MCJ243(*nhl-2 (ma371 cdb115 cdb120)* III), PQ583(*alg-2(ap431[3xflag::mkate2::alg-2]) II; alg-1(ap423[3xflag::gfp::alg-1]) X*). Some *C. elegans* strains used in this study were obtained from the *C. elegans* Genetics Center (USA), which is funded by the NIH Office of Research Infrastructure Programs (P40 OD010440). Strains were maintained on *Escherichia coli* OP50 and grown at 20 °C, unless different temperatures were required. RNAi knockdown was performed as previously described (72).

### *lin-28* 3’ UTR reporter

The *col-10* promoter and mCherry-H2B were amplified via PCR from worm genomic DNA, and the PEST domain, wild-type and mutated *lin-28* 3’ UTR were synthesized as a gene fragment (IDT gBlock). The wild-type and NHL-2 putative minding site mutations *lin-28* 3’ UTR sequences had three nucleotides of the *lin-4* miRNA seed sequences removed to inhibit action of this pathway (35). DNA fragments were assembled using the NEBuilder HiFi DNA Assembly Kit (New England Biolabs (NEB)) using the pGEM-T easy plasmid backbone. Constructs were injected as an extra chromosomal array at 1 ng/ul with *rol-6* marker into wild-type young adult hermaphrodite worms. All constructs were confirmed by sequencing prior to injection.

### Genome editing

Guide RNAs were designed in close proximity to the edit site, while favoring both high GC content and CRISPRscan scores (73,74). Alt-R crRNAs (IDT) were pre-annealed with tracrRNA at 10μM by heating to 95 °C for 5 min in IDT duplex buffer, then cooled for 5 min at room temperature. The injection mixes included 2μM Cas9 (IDT), 4μM total annealed guide RNAs (gKM39, gKM40), and a repair template with 35-nt homology arms (3). A *dpy-10* guide RNA (gKM3) and *dpy-10* repair template (oKM38) were used as markers of CRISPR efficiency in the injection mixtures (4). Two rounds of CRISPR were performed for *nhl-2* mutation. In the first round, injections were performed in the VT3554 (*gfp::3xflag::nhl-2*) strain to replace 211bp of *nhl-*2 sequence with a cleavage site for gKM3; the injection mix included gKM3, oKM38, gKM39, gKM40, and an ssDNA repair template (oKM259 at 120ng/μl). In the second round, this site was edited again with gKM3, oKM38, and a double-stranded repair template (dKM11 at 220ng/μl) to generate the desired *nhl-2* mutation.

### Microscopy

*C. elegans* was immobilized using 0.1% tetramisole (Sigma), placed on 2% agarose pads, and imaged using an Olympus 1X81/1X2-UCB fluorescence microscope (Olympus, Tokyo, Japan).

### Co-immunoprecipitation (Co-IP) and quantitative real-time PCR (qPCR)

RNA was extracted from total cell lysates directly from beads after IP using TRIzol RNA Miniprep Plus (Zymo Research). First, 300 µL of TRIzol was added to the RNA/protein sample and incubated at RT for 10 min. An equal volume of ethanol (95-100%) was added to the sample, and the mixture was pipetted into Zymo-Spin™ IIICG Column 2 in a collection tube. The samples were centrifuged at 16,000 *g* for 1 min at room temperature, and the flow-through was discarded. RNA Wash Buffer (400 µL) was then added to the column and centrifuged at 16,000 *g* for 1 min. The flow-through was discarded and the wash step was repeated. RNA Wash Buffer (700 µL) was added to the column and centrifuged for 2 min at 16,000 *g* to ensure complete removal of the wash buffer. The column was then carefully transferred to an RNase-free tube. To elute RNA, 20 µl of Dnase/RNase-Free Water was added directly to the column matrix, centrifuged at 16,000 *g*, and immediately stored frozen at-80°C. cDNA synthesis was performed using the ProtoScript® First Strand cDNA Synthesis Kit (NEB). One microliter of oligo d(T)23VN primer (50 μM) (NEB) was added to 200 ng/µL of RNA (170 ng worm RNA + 35 ng yeast RNA) or 500 ng/µL of total RNA extraction with 1µl 10 mM dNTP and RNAase-free water to reach 13 µL. The mixture was heated to 65 °C for 5 min and incubated on ice for at least 1 min. Then, 4 µl 5X First-Strand Buffer, 1 µl 0.1 M DTT, 1 µl RNaseOUT™ Recombinant RNase Inhibitor, and 1 µl of SuperScript™ III RT (200 units/µL) were added, followed by incubation at 55°C for 60 min and 70°C for 15 min to inactivate the reaction by heating then stored at −80 °C. qRT-PCR was performed using the Power SYBR Green Master Mix (Applied Biosystems). Each sample contained 10 µL of SYBR Green Mix, 0.6 µl of each primer (10 µM), 1 µL of cDNA and 7.8 µl of RNAse-free water. The samples were then run in triplicate. Relative abundance was calculated using the ΔΔCt method (Laviak and Schmittgen, 2001), using *act-1* as the internal control for total RNA extraction and yeast 18s for IP RNA extraction.

### Protein cloning, overexpression and purification

Protein expression constructs were constructed using the *E. coli* BL21 (DE3) expression system with ceNHL-2 NHL domain constructs (amino acid residues 729-1010) cloned into pGEX-6P-1 for GST-tagged protein expression. Rosetta 2 NEB 5-alpha competent *E. coli* (NEB) was used to clone the expression constructs. Overexpression and purification of GST-NHL protein by glutathione affinity chromatography have been reported for a previous ceNHL-2 NHL construct (Davis, 2018).

For crystallization, the GST tag was cut off using HRV 3C protease during an overnight dialysis step at 4°C, cleaving at a specific site between GST and NHL-2. Following cleavage, the NHL domain was subjected to size-exclusion chromatography on HiLoad 16/600 Superdex 75 (GE Healthcare) to purify the NHL domain from GST and bacterial proteins. The purity of the cleaved proteins was confirmed using SDS-PAGE.

For RNA-binding studies, the proteins were retained as GST-tagged constructs. Here, it should be noted that Mut[1-5], Mut[1-3] and Mut[4-5] mutations were made in an NHL construct encoding amino acids 729-1031 and compared to wild-type protein of the same length. Mutant constructs were constructed using a Q5 site-directed mutagenesis kit from NEB. To introduce specific mutations into the constructs, primers for site-directed mutagenesis were designed using the NEB web tool NEBasechange, and the recommended protocol from NEB was followed. Sequences of the mutant constructs were confirmed by Sanger sequencing, performed using Monash Micromon Genomics (Monash University).

### Crystallisation and Structure Determination

Initial small crystals were formed at 4 °C in 3.5 M sodium formate (Condition X of Index Screen; Hampton Research) in a sitting drop format as part of a robot-controlled screen in 96 well plates (200 nL drop size, 1:1 ratio). Larger protein crystals (0.7-1 mm) were produced in a range of conditions from to 3-3.5 M sodium formate, 0.1 M HEPES pH 6.8, in hanging drop format with a protein concentration of 5-10 mg/ml. (4 °C, 2 µL drop size, and protein-to-precipitant ratio of 1:1). These crystals were taken to the Australian Synchrotron MX2 beamline, where they were diffracted 1.4 Å resolution.

The crystals were cryoprotected in the mother liquor supplemented with 15% (v/v) glycerol and flash-cooled in liquid nitrogen. Diffraction images were collected at the Australian Synchrotron MX2 high-throughput protein crystallography beamline with an EIGER detector using the BLU-ICE acquisition software (75). The diffraction images were indexed using IMOSFLM39 and scaled using AIMLESS40 from the CCP4 suite (76). Phases were generated via molecular replacement (using PHASER) with one chain of drBRAT NHL as the search model (PDB ID: 4ZLR) (77). Subsequent model refinement was conducted using PHENIX43, model building was performed using COOT (78) and the data are available under the accession Number PDBID: 8SUC.

### Protein binding affinity measurements

Proteins were produced as described above and used in the binding studies prior to GST cleavage. Protein concentrations were measured using a Nanodrop 2000 (Thermo Scientific). RNA (17-mer and 10-mer U-rich nucleotides) with 5-end 6-FAM was purchased from Integrated DNA Technologies. Fluorescence anisotropy (FA) assays were performed by adding the protein (1 nM to 155 µM) and 5’ -FAM labelled RNA (1 nM) in 25 mM KCl, 20 mM HEPES KOH pH 7.4, 2 mM DTT, 2 mM MgCl2, 4% Glycerol to a final volume of 150 µM. Plates were placed in a PHERAstar plate reader (BMG Labtech) to measure the fluorescence polarization. Following the measurements on the Pherastar, the data were analyzed using MARS analysis software (BMG LABTECH, Germany). The average fluorescence anisotropy of the RNA alone was subtracted from the protein samples, which were then analyzed using GraphPad Prism (using a 1:1 specific site model) to determine the binding affinity of the protein to the RNA.

### Alphafold-multimer

To predict the structure of the heterodimer between CGH-1 and IFET-1, Alphfold2 (79) was run using default parameters in COSMIC².

## Acknowledgements.

We thank members of the Boag and Wilce laboratories for general discussions. Some of this work has been performed in the Monash Micromon core facility, we thank them for their support.

## Author Contributions

Conceptualization: PRB. Methodology and Investigation: NS, RNC, ALG, US, JWA. Funding acquisition: JMC, KM, JAW, PRB. Project Administration: PRB Supervision: JMC, MCJW, KM, JAW, PRB. Writing – original draft: PRB. All authors read and approved the manuscript.

---

**Supplemental Figure 1.**
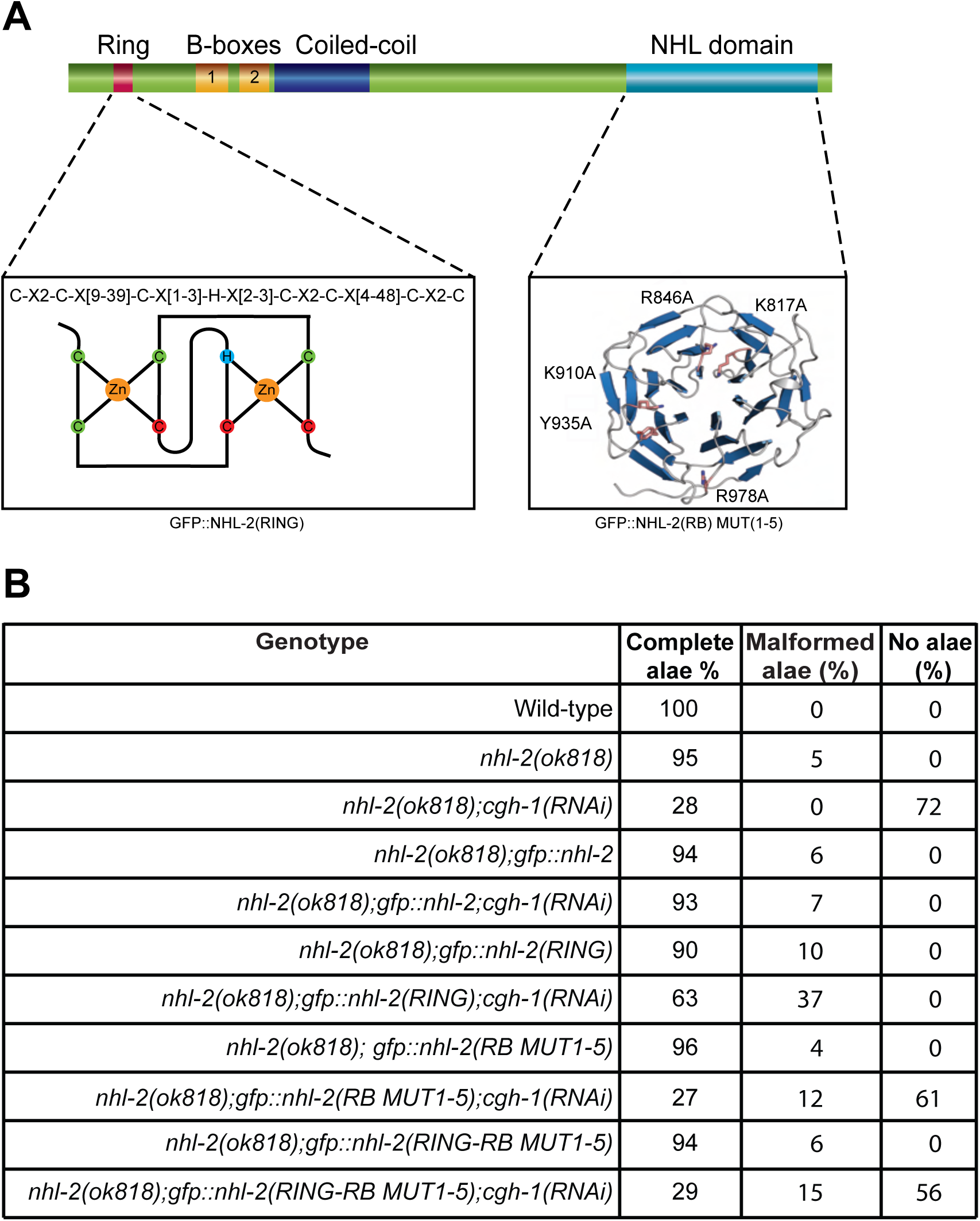
Analysis of genetic interactions of *nhl-2(ΔRING)* and *nhl-2(RB MUT1-5)* mutants. **A)** Illustration of the mutations made to the RING and NHL domains. The RING domain was rendered non-functional RING domain by replacing Cys77, Cys82, and Cys85 with alanine (red circles). RNA-binding activity was inhibited by replacing K817, R846, K910, Y935, and R978 with alanine). **B)** Quantification of alae defects in *nhl-2(RING)* and *nhl-2(RB MUT1-5)* worms with and without *cgh-1(RNAi)*.

**Supplemental Figure 2.**
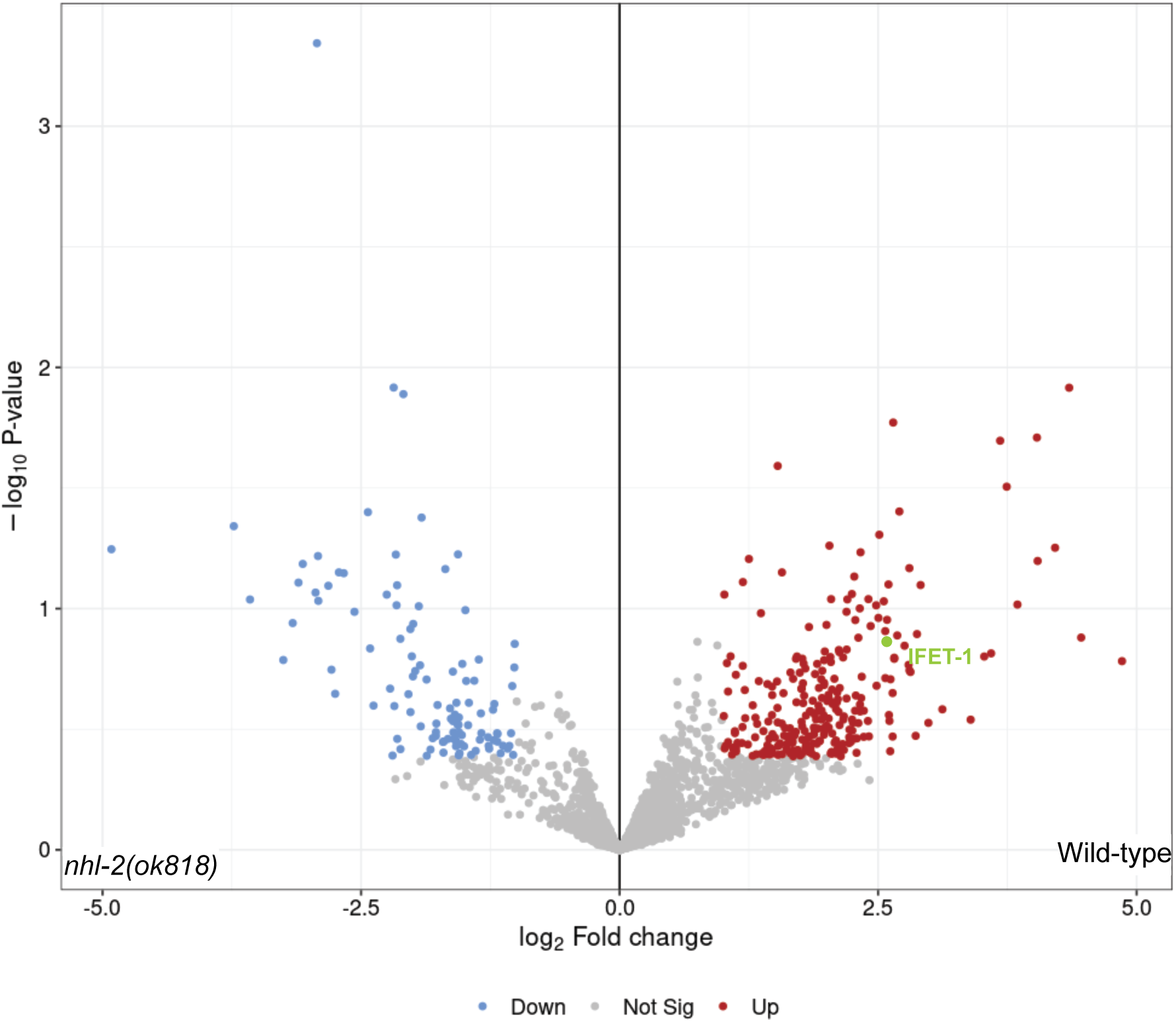
Volcano plot protein enrichment from mass spectrometry analysis of ALG-1/ALG-2 immunoprecipitations. IFET-1 is significantly enriched in wild-type compared to *nhl-2(ok818)* worms.

